# The Reissner fiber under tension in vivo shows dynamic interaction with ciliated cells contacting the cerebrospinal fluid

**DOI:** 10.1101/2023.02.22.529498

**Authors:** Celine Bellegarda, Guillaume Zavard, Lionel Moisan, Ryan S. Gray, Françoise Brochard-Wyart, Jean-François Joanny, Yasmine Cantaut-Belarif, Claire Wyart

## Abstract

The Reissner fiber (RF) is an acellular thread positioned in the midline of the central canal that aggregates thanks to the beating of numerous cilia from ependymal radial glial cells (ERGs) generating flow in the central canal of the spinal cord. RF together with cerebrospinal fluid (CSF)-contacting neurons (CSF-cNs) forms an axial sensory system detecting curvature. How RF, CSF-cNs and the multitude of motile cilia from ERGs interact *in vivo* appears critical for maintenance of RF and sensory functions of CSF-cNs to keep a straight body axis but is not well-understood. Using *in vivo* imaging in larval zebrafish, we show that RF is under tension and resonates dorsoventrally. Focal RF ablations trigger retraction and relaxation of the fiber cut ends, with larger retraction speeds for rostral ablations. We built a mechanical model that estimates RF stress diffusion coefficient at 4 mm^2^ / s and reveals that tension builds up rostrally along the fiber. After RF ablation, CSF-cN spontaneous activity decreased and ciliary motility changed, suggesting physical interactions between RF and cilia projecting into the central canal. We observed that motile cilia were caudally-tilted and frequently interacted with RF. We propose that the numerous ependymal motile monocilia contribute to RF heterogenous tension via weak interactions. Our work demonstrates that under tension, the Reissner fiber dynamically interacts with motile cilia generating CSF flow and spinal sensory neurons.

## INTRODUCTION

The cerebrospinal fluid (CSF) is secreted by the choroid plexuses and fills the intracerebral ventricles, spinal and brain subarachnoid spaces and the central canal of the spinal cord (Lun *et al*., 2015; Battal *et al*., 2011). In the last decade, the physico-chemical properties of the CSF have been shown to impact the development of the nervous system by influencing neurogenesis (Lehtinen *et al*., 2011), neuronal migration (Paul *et al*., 2017) as well as morphogenesis (Bearce & Grimes, 2019). In zebrafish, the circulation of CSF modulates the geometry of the body axis during embryogenesis (Zhang *et al*., 2018) and of the spine curvature in juvenile/adult zebrafish (Grimes *et al*., 2016). Genetic investigations in zebrafish revealed that straight body axis formation and spine organogenesis rely on the expression of urotensin neuropeptides in the spinal cord by ciliated sensory neurons surrounding the central canal, referred to as CSF-contacting neurons (CSF-cNs) (Quan *et al*., 2015; Zhang *et al*., 2018; Bearce *et al*., 2022; Gaillard *et al*., 2022).

Spinal CSF-cNs were identified by Kolmer and Agduhr in over a hundred vertebrate species (Kolmer, 1921; Agduhr, 1922; Vigh & Vigh-Teichmann, 1998) as GABAergic sensory neurons that project an apical extension into the lumen of the central canal and extend an ascending axon into the spinal cord. The apical extension comprises one motile cilium and a brush of microvilli that bathe in the CSF (Vigh & Vigh-Teichmann, 1998). During development, CSF-cNs differentiate into dorsolateral and ventral populations that are distinguished by distinct developmental origins (Park *et al*., 2004; Huang *et al*., 2012; Petracca *et al*., 2016). These two populations differ by the morphology of their ciliated apical extension and axonal projections, expression of peptides (Quan *et al*., 2015; Djenoune *et al*., 2017; Desban *et al*., 2019; Prendergast *et al*., 2019) as well as the neuronal targets contacted by the CSF-cN axons (Djenoune *et al*., 2017; Desban *et al*., 2019; Orts-Del’Immagine *et al*., 2014).

CSF-cNs express the urotensin-related peptides under the control of the Reissner Fiber (RF) (Quan *et al*., 2015;Zhang *et al*., 2018; Cantaut-Belarif *et al*., 2020; Lu et al., 2020), an acellular thread bathing in CSF within the lumen of the central canal and in close vicinity to the apical extension of CSF-cNs. Thanks to the beating of the numerous motile monocilia from ependymal radial glia, RF forms during development from the aggregation of the monomer SCO-spondin, a very large glycoprotein (Sterba et al., 1982; Cantaut-Belarif *et al*., 2018; Troutwine *et al*., 2020). SCO-spondin is initially secreted into the CSF by cells in the floor plate, the flexural organ and the subcommissural organ (SCO), and is later solely produced by the SCO (O. Meiniel & A. Meiniel, 2007; Gobron *et al*., 2000). In larval zebrafish, RF diameter is approximately 200 nm (Orts-Del’Immagine *et al*., 2020) and the RF displays slow, continual rostrocaudal movement (Troutwine *et al*., 2020). In mutant embryos for *scospondin* in which the RF does not form, the body axis is curled down (Cantaut-Belarif *et al*., 2018). Furthermore, in *scospondin* mutants in which the RF forms in the embryo but is not maintained in juvenile stages (Troutwine *et al*., 2020, Rose *et al*., 2020), the body axis becomes straight during embryogenesis but later on, the spine undergoes 3D torsion at the juvenile stage, a hallmark of idiopathic scoliosis. Peptides from the family of urotensin (Urp1, Urp2) and their receptor Uts2r3 play a major role in straightening the body axis throughout life (Bearce *et al*., 2022; Gaillard *et al*., 2022). Remarkably, urotensin signaling is relevant in human patients with adolescent idiopathic scoliosis (Dai *et al*., 2021; Xie *et al., in press*).

We previously showed that CSF-cNs sense zebrafish body curvature *in vivo* (Böhm *et al*., 2016). Accordingly, CSF-cNs can respond *in vitro* to pressure-application in an open book preparation of the lamprey spinal cord (Jalalvand *et al*., 2016) as well as to mechanical stimulation applied to CSF-cN cell membranes in primary cultures (Sternberg *et al*., 2018). In zebrafish, CSF-cNs sense spinal curvature *in vivo* only on the concave side (Böhm *et al*.,2016), a process that requires intact Pkd2l1 channels. This process relies on a modulation of the opening probability of the mechanosensory channel Pkd2l1 (Sternberg *et al*., 2018) for CSF-cNs on the concave side, ipsilateral to where skeletal muscles contract (Böhm *et al*., 2016). In zebrafish lacking RF, both the CSF flow (Cantaut-Belarif *et al*., 2018) and Pkd2l1 channel activity (Sternberg *et al*., 2018) are unaffected, yet CSF-cN response to spinal curvature was reduced (Orts-Del’Immagine *et al*., 2020). This observation suggests that an intact RF amplifies the mechanosensitivity of CSF-cNs, either by RF contacting the ciliated CSF-cN apical extension or by enhancing the gradient of CSF flow that CSF-cNs are subject to during spinal bending. Previous investigations on the structure of RF and its interaction with CSF-cNs were performed in fixed tissues (using 4% paraformaldehyde; see Cantaut-Belarif *et al*., 2018; Orts-Del’Immagine *et al*., 2020), which leads to deformation of the central canal that can alter the relative distance of the RF to CSF-cNs (Orts-Del’Immagine *et al*., 2020; Thouvenin *et al*., 2020). It is therefore not yet clear how CSF-cNs activity is affected by interacting with the RF *in vivo*.

In this study, we took advantage of the transparency of transgenic zebrafish larvae to investigate *in vivo* the dynamic properties of the RF and its interaction with ciliated cells contacting the CSF in the central canal. By developing a tracking method to identify the dorsoventral position of the RF along the rostrocaudal axis, we discovered that at the larval stage, the RF is under tension and undergoes dynamic spontaneous oscillatory activity *in vivo* over 100-200 nm in the dorsoventral axis, with largest amplitudes of oscillations in the middle portion of the fish. Interestingly, the focal ablation of the RF using a pulsed laser led to an initial fast retraction of the RF with maximal retraction speeds reaching up to 700 μm/s when the ablation was performed rostrally. After an initial retraction, relaxation of the RF was sometimes accompanied by a retention of the cut end in the central canal with an observable deformation of the fiber when it seemed to interact with other cellular components in the CSF. We built a mechanical model of the RF to estimate from the relaxation kinetics its elastic properties, including its mechanical diffusion coefficient, characteristic time, and retraction ratio. Our model revealed a heterogeneous tension along the RF, larger on the rostral portion, which is not likely explained by CSF flow. In order to investigate whether motile cilia, including the most numerous ones from ependymal radial glial cells (ERGs) and the sparse ones from CSF-cNs, were interacting with the RF, we performed acute focal ablations of the RF using pulsed lasers. We found that RF ablation decreased spontaneous calcium activity in CSF-cNs. We observed as well frequent interactions between the RF and the tip of beating monocilia. Accordingly, we found that the frequency of ciliary beating was often affected by RF ablation. Together with our observations that the polarity of cilia beating in the sagittal plane was biased caudally, our findings suggest that caudally-oriented beating of monocilia may contribute to the generation of an heterogeneous tension via weak interactions between motile cilia and the fiber. Altogether, our observations indicate that the RF is a dynamic structure under tension in the CSF that interacts with CSF-cNs as well as with the motile monocilia in the central canal. The dynamic RF subsequently promotes activity in a subset of sensory neurons at rest, while the implications of the interactions between RF and long motile cilia from ependymal radial glia remain to be investigated.

## RESULTS

### 1/ The Reissner fiber under tension spontaneously oscillates *in vivo*

To investigate the dynamical properties of the Reissner fiber, we performed high-speed imaging of the central canal in the transgenic reporter knock-in zebrafish line *Tg(sspo:sspo-GFP*) (Troutwine *et al*., 2020), in which the fiber is GFP-tagged (Figure 1). In 3 days post fertilization (dpf) *Tg(sspo:sspo-GFP*) larvae after paralysis (see Methods), we observed that the Reissner fiber was straight and taut in the sagittal plane and rapidly changed position along the dorsoventral axis over time in the central canal (Figure 1A, B). These observations indicate that the fiber is under tension *in vivo*. In contrast, the Reissner fiber of larvae after fixation was slack and stationary (Figure 1C; see also Movies 1 in live larvae, 2 after fixation). We developed a script to track the position of the fiber in the dorsoventral axis (Figure 1D; see also Methods) as a function of its position in the rostrocaudal axis, discretized in 2 μm bins (Figure 1E1, 1E2). We estimated that the dorsoventral displacement of the fiber from its mean position in the central canal (on average median ± standard deviation for all values provided hereafter: 74 nm ± 68 nm) was significantly larger in paralyzed living larvae than in euthanized larvae after fixation (on average: 32 nm ± 39 nm; p<1×10^-4^; unpaired two-tailed t-test; Figure 1F1), whose perceived displacement may be due to noise in our imaging setup and artifacts of detection of the center position of the fiber. The amplitude of the Reissner fiber dorsoventral displacement was largest in the middle portion (125 nm ± 108 nm), followed by the fiber displacement on the rostral side (100 nm ± 103 nm; p<1×10^-4^; Tukey’s HSD Test for multiple comparisons; Figure 1F1). The median of the amplitudes of Reissner fiber displacement in the caudal end (70 nm ± 68 nm; p<1×10^-4^; Tukey’s HSD Test for multiple comparisons; Figure 1F2) was the closest to the fiber displacement of fixed larvae, possibly partly reflecting that the fiber is anchored on both the rostral and caudal ends of the fish. Recordings from rostral, middle and caudal portions of different fish showed a greater variability in dorsoventral displacement from the mean position in the middle portion of the fish than on the rostral and caudal sides (Figure 1F3). Performing a spatial principal component analysis over all pixels in the movie revealed that the first three components after dimensionality reduction represented the dorsoventral translation of the fiber moving either dorsally (Figure 1G top panel; see also Movie 3) or ventrally (Figure 1G middle panel; see also Movie 3). These observations uncover that the Reissner fiber under tension *in vivo* demonstrates dynamic dorsoventral oscillations with graded amplitudes of oscillations along the rostrocaudal axis.

**Figure 1.**
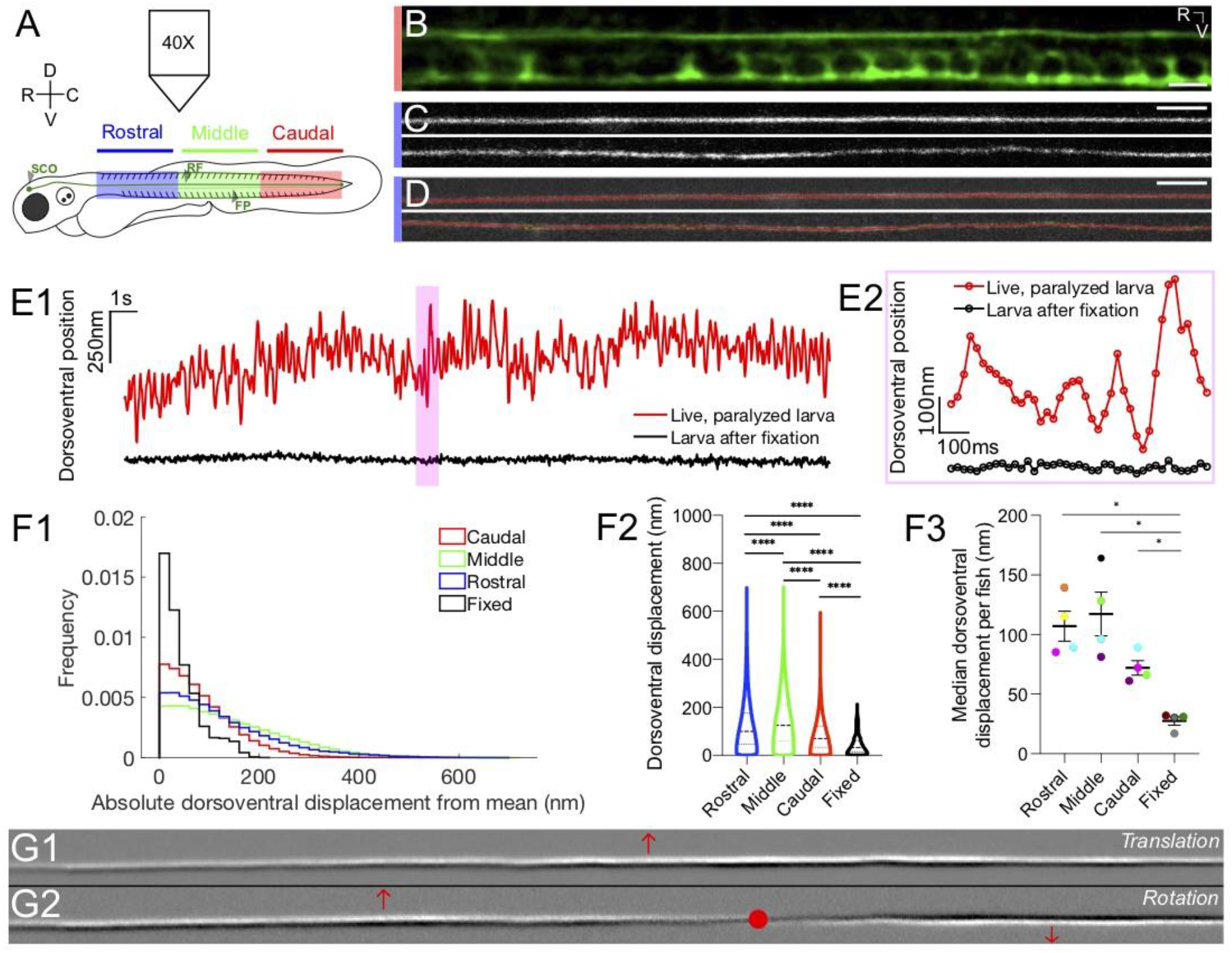
The Reissner fiber under tension exhibits spontaneously dynamic behavior over the dorsoventral axis in the central canal. (A) Spinning disk confocal microscopy setup using a 40X objective with 3 dpf *Tg*(*sspo:sspo-GFP*) zebrafish larvae for live imaging. Schema of zebrafish larva designates Rostral (blue), Middle (green) and Caudal (red) sections, corresponding to somites 1-10, 11-20, 21-30, respectively. (B) Immunohistochemistry with anti-GFP antibodies in 3 dpf *Tg*(*sspo:sspo-GFP*) larva after fixation shows the Reissner fiber (RF) with the floor plate (FP) visible within the caudal somites of the spinal cord. (C) Live imaging snapshot of the RF in the rostral somites of a 3 dpf *Tg*(*sspo:sspo-GFP*) paralyzed, living larva (top) and larva after fixation (bottom). (D) Example tracking of continuous motion of the RF through the development of a script to model its movements in the dorsoventral axis. (E1) Example trace of the change in dorsoventral position of the RF in one paralyzed, living larva (red) and another euthanized larva after fixation (black) over a 25 s-long timelapse acquired at 40 Hz. Data was discretized in 2-μm bins along the rostrocaudal axis before plotting. (E2) Zoomed-in display of the highlighted area marked on E1, showing a trace of the dorsoventral position of the RF over 1 s for both the paralyzed, living larva and the euthanized larva after fixation, respectively, with circles indicating the sampling points. (F1) Displacement in the dorsoventral axis for paralyzed larvae (4 Rostral, 4 Middle, 4 Caudal recordings) is significantly larger than that of fixed larvae (4 recordings) (on average median ± standard deviation provided hereafter: in paralyzed living larvae = 74 nm ± 68 nm versus in fixed larvae = 32 nm ± 39 nm; unpaired two-tailed t-test: p < 10^-4^). The displacement was calculated from data that was discretized in 2-μm bins along the rostrocaudal axis. (F2) Dorsoventral displacement of the RF is significantly different among rostral, middle and caudal segments of paralyzed larvae (on average median ± standard deviation in rostral somites = 100 nm ± 103 nm versus in middle somites = 125 nm ± 108 nm versus in caudal somites = 70 nm ± 68 nm versus in fixed larvae = 32 nm ± 3 9 nm; Tukey’s HSD Test for multiple comparisons: p < 10^-4^). (F3) Median dorsoventral displacement of the RF from the mean position per fish, with each color representing a different fish (on average mean of median dorsoventral displacement ± standard deviation in rostral somites = 107 nm ± 25 nm versus in middle somites = 117 nm ± 37 nm versus in caudal somites = 72 nm ± 12 nm versus in fixed larvae = 28 nm ± 7 nm; Tukey’s HSD Test for multiple comparisons: p < 0.05). (G1-G2) A principal component analysis was computed on one image sequence of one fish (with each image corresponding to one observation) to understand the most significant movements of the fiber. (G1), (G2) respectively represent the first and second component of the PCA, which can be interpreted as temporal gradients (white, intensity increases and black, intensity decreases). The first component corresponds to a dorso-ventral translation (the fiber borders are black on the ventral side, white on the dorsal side) and the second to a small local rotation around point I (in red). These two components account respectively for 21.9% (G1) and 3.4% (G2) of the total temporal variability in the movie. * p < 0.05, **** p < 10^-4^. Scale bar is 10 μm (B, C, D) and 20 μm (G).

### 2/ The Reissner fiber enhances spontaneous calcium activity in cerebrospinal fluid-contacting neurons

Given that the fiber in paralyzed, living larvae exhibits spontaneous dorsoventral movements, we investigated whether these oscillations contribute to the spontaneous calcium activity of the CSF-cNs (Figure 2). We performed acute two-photon ablations of the Reissner fiber and tested whether it impacted the spontaneous calcium activity of CSF-cNs in triple transgenic *Tg*(*sspo:sspo-GFP;pkd2l1:tagRFP;pkd2l1:GCaMP5G*) paralyzed larvae (Figure 2A1, 2A2; see also Movie 4 before RF ablation, Movie 5 after RF ablation and Methods). To record the spontaneous activity of CSF-cNs before and after RF photoablation, we monitored calcium transients of ventral CSF-cNs located in the sagittal plane of RF (Figure 2B). Overall, fewer cells were active after RF photoablation (on average 11% compared to 28%; p < 0.05; paired two-tailed t-test; Figure 2C), and calcium activity decreased by 45% on average across larvae (Figure 2D) with a subset of ventral CSF-cNs showing decreased activity after RF photoablation. The number of calcium events occurring within those cells decreased after ablation (Figure 2E, on average 0.94 events / min before and 0.87 events/min after in active cells; p < 0.005; paired twotailed t-test). Altogether, our results indicate that the presence of an intact Reissner fiber in the central canal enhances the spontaneous calcium activity of ventral CSF-cNs.

**Figure 2.**
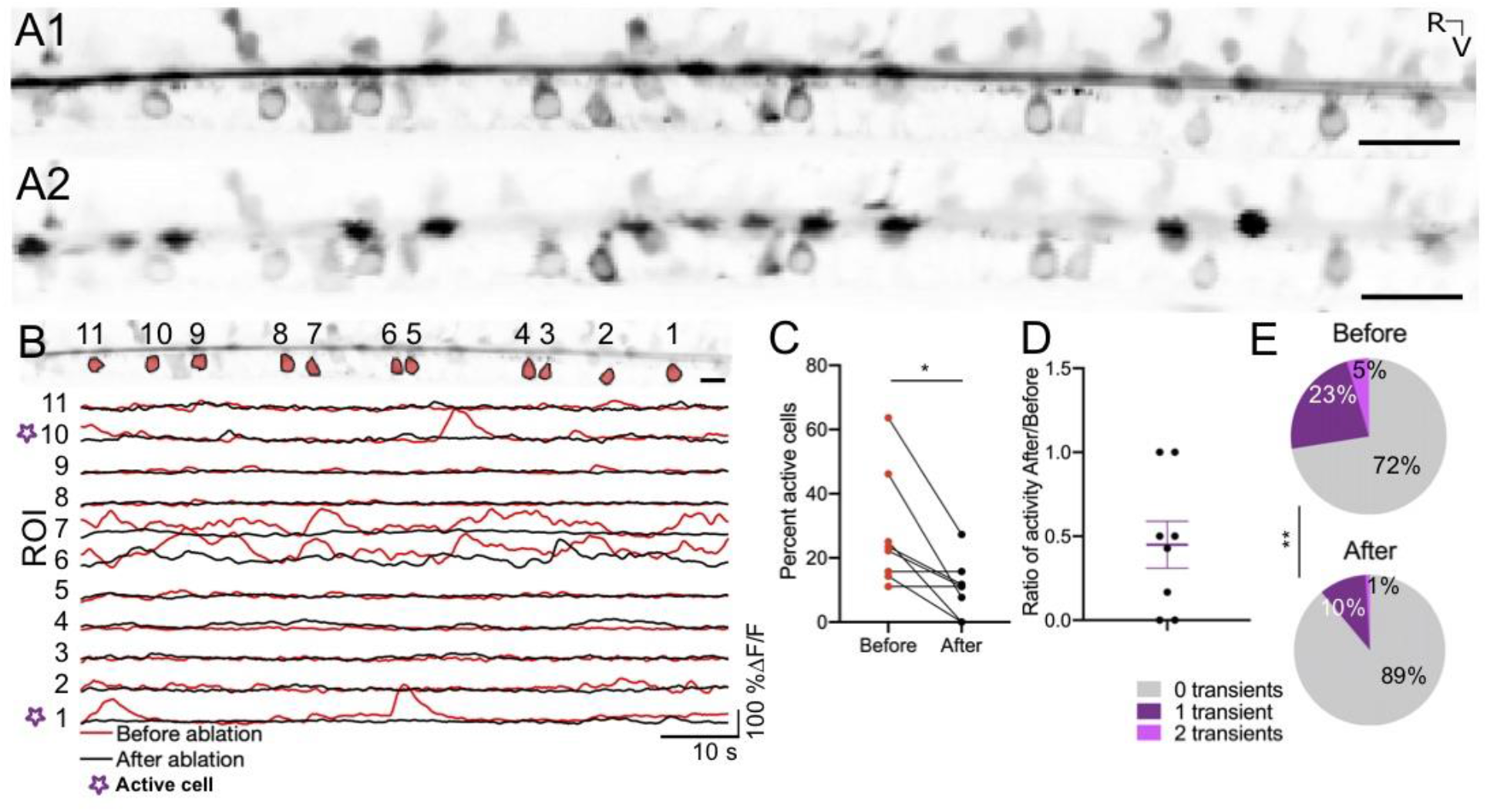
The Reissner fiber enhances spontaneous calcium activity in ventral CSF-cNs. (A1) Time-series standard deviation projection in the sagittal plane from 2 photon laser scanning microscope showing the signal from the Reissner fiber and CSF-cNs in the central canal of 3 dpf *Tg*(*sspo:sspo-GFP;pkd2l1:tagRFP;pkd2l1:GCaMP5G*) zebrafish larva before RF photoablation. (A2) Time-series standard deviation projection in the sagittal plane from 2 photon laser scanning microscope showing the signal from CSF-cNs after RF ablation performed by spiral scanning photoablation with an infrared pulsed laser tuned at 800 nm over 0.5 μm on the RF (see Methods). (B) ROI selection for ventral CSF-cNs to analyze activity before and after RF photoablation within the same cells (top). Example calcium activity traces normalized to baseline for each of the ROIs before (red) and after (black) RF photoablation over 75 s imaged at 3.45 Hz (see Methods). (C) Percentage of active ventral CSF-cNs (active is defined as having at least 1 calcium transient for the recording) before and after RF photoablation (109 cells total from 8 fish from 2 independent clutches; mean percent active before ablation = 27.72% ± 6.34% versus mean percent active after ablation = 10.59% ± 3.1%; paired two-tailed t-test: p < 0.05). (D) The ratio of active ventral CSF-cNs after RF photoablation to those active before RF photoablation, illustrating on average, a fraction (on average ± SEM: 45% ± 14%) of active ventral CSF-cNs before photoablation remain active after RF photoablation. The purple lines on the graph represent the mean and the error bars indicate the SEM. (E) Pie charts illustrating the number of events per ventral CSF-cN before and after RF photoablation (mean number of events in active cells before RF photoablation = 0.94 events / min versus mean number of events in active cells after RF photoablation = 0.87 events / min; paired two-tailed t-test: p < 0.005). *p < 0.05, ** p < 0.005 Scale bar is 20 μm (A1, A2), 10 μm (B).

### 3/ Estimation of the elastic properties of the Reissner fiber from acute ablation

To gain a deeper understanding of the elastic properties of the RF, we explored its response to acute photoablation. We performed RF photoablations via a UV-pulsed laser system in triple transgenic *Tg*(*sspo:sspo-GFP;pkd2l1:tagRFP;pkd2l1:GCaMP5G*) zebrafish larvae (N=74 fish). We then tracked the relaxation dynamics of the two cut ends of the fiber (Figure 3; see Movies 6-8; see also Methods). We observed diverse kinetics and retraction patterns upon photoablation. The majority of fiber retractions (88% of 148 total retractions, Figures 3A1, 3A2) demonstrated complete fiber retraction out of the 97 μm wide field of view in less than 500 milliseconds (initial retraction speed of on average ~ 328 μm/s) with 96% of fibers remaining straight as the cut ends of the fiber retracted to the rostral and caudal ends of the larvae. However, in a few cases (12% of 148 total retractions, Figure 3B) we observed the two cut ends of the fiber relaxed very slowly (initial retraction speed of on average ~ 50 μm/s), often with cut ends remaining still in the field of view for over 20 s (Figures 3C1, 3C2, 3C3). In a third of these relaxed cases, we could observe that slow-retracting ablated fibers displayed at the tip snake-like deformations during their relaxation (Figure 3B). As an analogy with the dynamic model of DNA molecules (Brochard-Wyart, 1995), we refer to the deformed section of the fiber as the “flower”, which suggests the fiber was no longer under tension (Figure 3B2). In this analogy, the flower contrasts with the straight “stem” section of the fiber that remained under tension (Figure 3B3).

To estimate the elastic properties of the fiber, in particular its mechanical diffusion coefficient, we developed a simple model for the ablation of an elastic fiber inspired by the dynamic model for DNA molecules (Brochard-Wyart, 1995). The Reissner fiber can be seen as an elastic rod-like polymer in the central canal with a radius r_f_ of ~100 nm (Orts-Del’Immagine *et al*., 2020). The relation between the pulling force F acting on the fiber and its deformation can be described as:

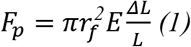

with E describing the elastic Young modulus, ΔL the elongation of the fiber stretched away from its original full length L when it is under tension *in vivo*.

When the fiber is cut, the deformation relaxes from the free end over a distance (x) e.g. “ the flower” while the rest of the fiber e.g. “the stem” remains under tension. The size of the flower is deduced from a balance between the pulling force (Equation 1) and the friction force F_v_ acting on the flower, which can be written as:

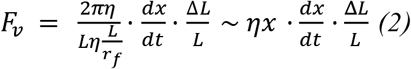

With x being the size of the relaxed fiber (flower), η the CSF viscosity and 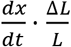 the retraction velocity that leads to friction.

The force balance *F_p_* = *F_v_* leads to a diffusion equation and therefore

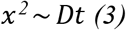

where D is given by 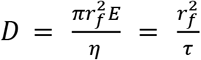 with the characteristic time *τ* for the fiber mechanical relaxation 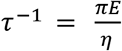.

Based on our model, the retraction distance upon ablation should increase as a function of √t. Because the deformation of the fiber in the flower is relaxed, the retraction X of the fiber given by the position of the free end is:

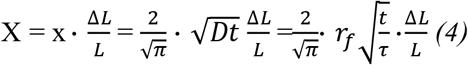

By plotting the retraction position as a function of √t (Figure 3C1), we found indeed as expected from our simple model a linear relationship (Figure 3C1). The slope of this relation defined by 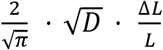 enabled us to group the fibers with fast retraction (slope = 161 μm/√s) and slow retraction (slope = 10 μm/√s) (Figure 3C1). To assess whether the retraction kinematics of the fiber differ along the rostrocaudal axis, we compared the kinematics of retraction after acute RF photoablations performed at different sites (using the same terminology “rostral”, “middle” and “caudal”, see Figure 1A). When ablations occurred in the rostral side, we only observed fast retraction kinetics (n = 105, Figure 3D). In contrast, ablations in the middle and caudal somites showed a slower relaxation speed (Figure 3D). The retraction speed for the rostral and caudal cut end of a given fiber were highly correlated in individual larvae despite the diversity of retraction patterns observed overall across fish (y = 0.9 x + 2; p < 1×10^-4^; R^2^=0.7; simple linear correlation; Figure 3D), indicating that the retraction kinematics after ablation reveals the inherent physical properties of the fiber and the tension applied on it that differs as a function of the rostrocaudal position.

From the slope 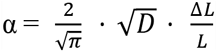 of the retraction distance as a function of 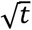, we can extract from the information on the change of RF length 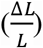 the mechanical diffusion coefficient D. In ablation cases with retention of the fiber that remained in the field of view, we estimated ΔL the change of length of the fiber as it stretches away by the distance between the retention position of the cut end once immobile and the ablation locus. ΔL was negatively correlated with the position of the ablation locus along the rostrocaudal axis (y = - 0.9 x + 41; p < 0.06; R^2^ = 0.2; simple linear regression; Figure 3E). Accordingly, maximum retraction speeds were greater in the rostral somites of the larvae (on average ± standard deviation for all values provided hereafter: 448 μm/s ± 168 μm/s) than those in the middle or caudal somites (253 μm/s ± 166 μm/s and 211 μm/s ± 165 μm/s, respectively; Tukey’s HSD Test for multiple comparisons; p < 10^-4^; Figure 3F).

In 13 cases of ablation on the caudal side, the fiber cut ends remained in the field of view during the 38 s - long recording and we could therefore measure ΔL as low as 10 μm. For rostral ablation, based on our field of view of 97 μm, we can estimate ΔL > 150 μm (rostral ablation) on each side. These values lead for a fiber length of approximately 3 mm long to ΔL / L_Rostral_ >~ 1/20 and ΔL / L_Middle/Caudal_ ≃ 1/300.

Using the slope α, we grouped the fibers with fast retraction (slope = 161 μm/√s) versus the fibers with slow retraction (slope = 10 μm/√s) (see Figure 3D). We found a remarkably-similar estimation of the mechanical diffusion coefficient for both group with D ≃ 5.6 mm^2^/s (for fibers ablated on the rostral side and showing fast retraction: slope = 0.161 mm/√s; ΔL/L_Rostral_ ≃ 1/20; D_Rostral_≃ 5.7 mm^2^/s, and for fibers ablated in the more caudal position and showing slow retraction: slope = 0.010 mm/√s; ΔL/L _Middle/Caudal_ ≃ 1/300; D_Middle/Caudal_ ≃ 5.2 mm^2^ / s). Consequently, the characteristic time for the fiber mechanical relaxation 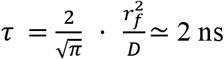. The total retraction time is 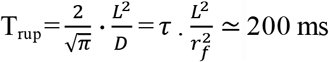 as observed experimentally.

The fit with experimental data shows that D is uniform along the fiber, and the faster retraction dynamics in the rostral than the caudal region demonstrates that the fiber tension increases along the rostrocaudal axis, from caudal to rostral. We can suggest two interpretations. The first one is the stretching of the fiber by frictional forces of the cerebrospinal fluid flowing from rostral to caudal with velocity U. At a distance *l* from the caudal end, the hydrodynamic pulling force 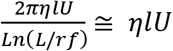, increasing from the caudal (1 = 0) to the rostral (1 = L), is balanced by the fiber tension 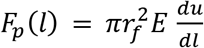, where 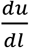is the fiber deformation.

It leads to an increase of the rostral deformation 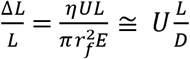. The second contribution may be due to the stretching by the cilia of ependymal radial glial cells (the most numerous) exerting a force *f_p_* on the fiber in the flow direction. If *v* is the linear density of the cilia-fiber links pulling with the force *f_p_*, the resulting pulling force at a distance l from the caudal end is 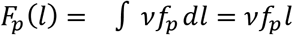. The pulling force increases from the caudal to the rostral, leading to a maximal rostral deformation 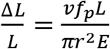. From the value of D, we can estimate the value of the fiber elastic modulus 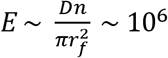 Pascal.

Overall, the Reissner fiber in larval zebrafish can be described as an elastic polymer that is maintained under tension in the CSF and exhibits a mechanical diffusion coefficient of about 4 mm^2^/s, a characteristic time in the order of 2 ns and an elastic modulus of 10^5^ Pascal – a value that would correspond to a low density gel rather than a rigid protein fiber.

**Figure 3.**
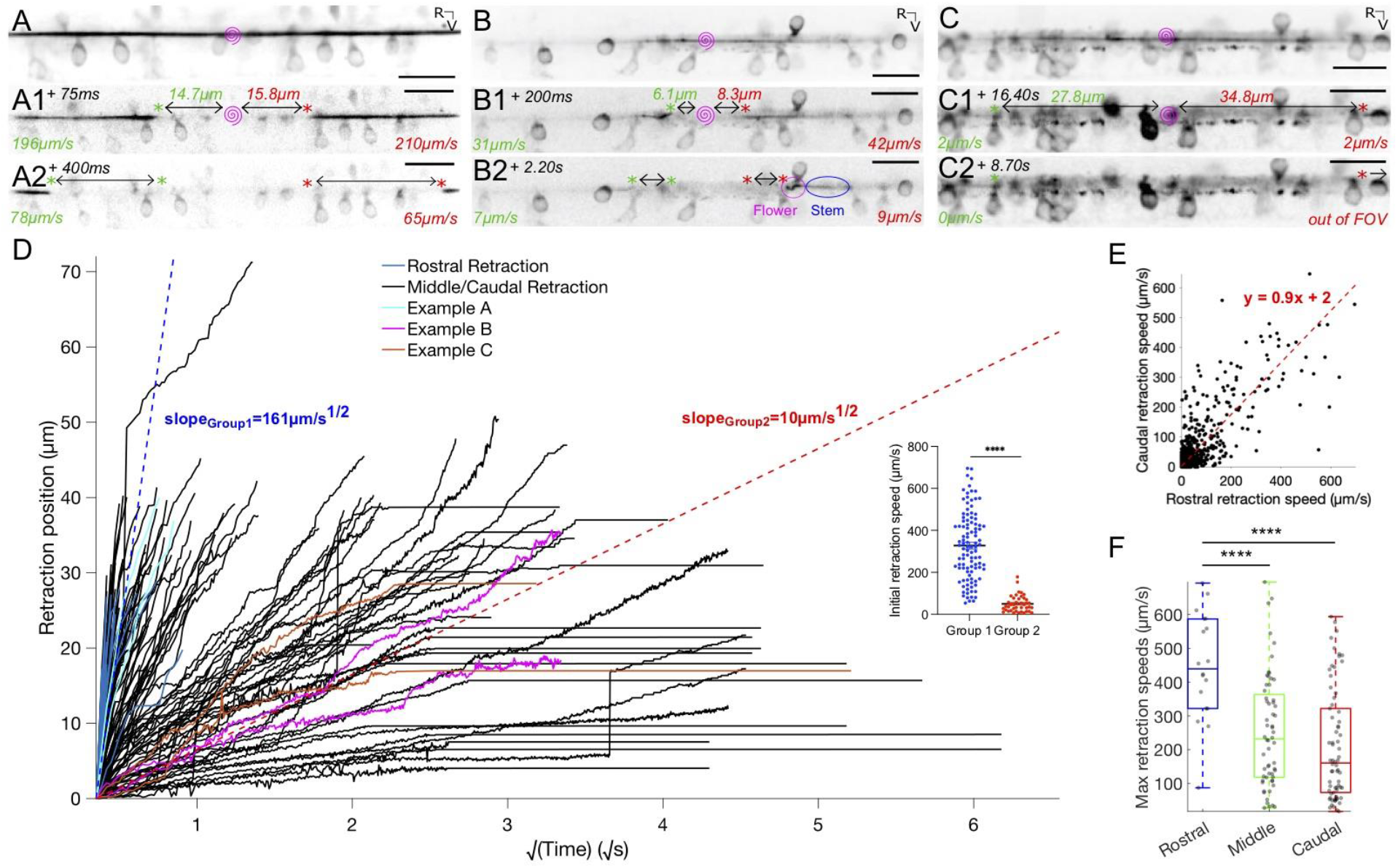
Reissner fiber photoablation reveals different modes of relaxation under tension. (A) Frame-by-frame instantaneous speed calculations for the rostral and caudal ends of the cut Reissner fiber during retraction after photoablation by UV pulsed laser in 3 dpf triple transgenic *Tg*(*sspo:sspo-GFP;pkd2l1:tagRFP;pkd2l1:GCaMP5G*) paralyzed larvae. Pink spiral indicates the site of RF ablation. Instantaneous speeds of the fiber for both rostral and caudal ends were calculated using the change in position of the fiber between two consecutive frames, divided by the exposure time (25 ms) and converted to μm/s. (A1, A2) specifically depicts an example of a fiber in the “rigid case”, where the fiber retracts straight on both ends without distortion. (B) Instantaneous speed calculations for both rostral and caudal ends of a cut fiber in the “relaxed case”, where there is distortion of the fiber (B1 left cut end, B2 right cut end) due to no more tension on one or both ends of the fiber. By analogy with the dynamics of DNA molecules, one can refer to the completely relaxed portion of the fiber as the “flower”, while the remaining taut portion of the fiber would be the “stem”. (C) Instantaneous speed calculations for both rostral and caudal ends of a cut fiber that remains stuck in the central canal, remaining in the field of view for about 25 s. (D) Retraction position plotted across the square root of time for all fish (N = 74 fish from 7 independent clutches), color-coded demonstrating the examples in (A-C), and indicating if a flower was seen in the 38 s-long recordings along the rostrocaudal axis: √D * (ΔL/L) is provided by the slopes of the two dotted red lines, indicating a fast retraction or a slower retraction of the rostral and caudal ends of the fiber in the central canal. Inset: initial retraction speed (the distance the cut fiber retracted in between 50 ms and 75 ms after photoablation, after UV laser artifacts) is larger in the fibers classified under the fast retraction group than those in the slow retraction group. (E) Rostral and caudal ends of the fiber in cases with and without flowers plotted against each other, demonstrating that the rostral and caudal end dynamics generally tend to mirror each other (74 fish). (F) Maximum retraction speed for the rostral and caudal ends of the ablated RF classified using the schema in Figure 1A, grouping the ablations that occurred in the rostral somites (Nb=b9 fish), middle somites (34 fish), and caudal somites (34 fish). Mean for ablation in rostral somites = 448 μm/s ± 168 μm/s versus in middle somites = 253 μm/s ± 166 μm/s versus in caudal somites = 211 μm/s ± 165 μm/s; p<1×10^-4^; Tukey’s HSD Test for multiple comparisons. ****p < 1×10^-4^ Scale bar is 10 μm (A, B).

### 4/ The Reissner fiber in the larva interacts with motile cilia along the central canal

The RF appears to directly interact with CSF-CN cilia in fixed tissues (Orts-Del’Immagine *et al*., 2020). Given that the RF displays diverse retraction patterns along the rostrocaudal axis after photoablation, we investigated whether the fiber may dynamically interact with cilia in the central canal in vivo (Figure 4). Ciliated cells in contact with the CSF include CSF-cNs which project a short motile monocilium (Böhm, U., Prendergast, A., Djenoune, L. *et al*., 2016) known to be about a hundred at this stage (Prendergast *et al., in press*), as well as ependymal radial glial cells, which project a longer monocilia (Borovina, A., Superina, S., Voskas, D. *et al*., 2010; Becker, C. & Becker, T., 2015) and are about hundreds of thousands at this stage. We asked whether the latter may interact with the RF to elicit the apparent friction and build a heterogenous tension along the fiber.

We investigated whether we could find evidence for an interaction between the RF and motile cilia using high-speed imaging in double transgenic *Tg(sspo:sspo-GFP;β-actin:Arl13b-GFP*) larvae, in which many cilia in the central canal are labeled with GFP (Figure 4B; see also Methods). Immunohistochemistry staining against glutamylated tubulin and GFP showed that motile cilia are densely packed along the dorsal and ventral walls of the central canal and are oriented with a caudal tilt (Figure 4A). In contrast to the embryo in which only ventral cilia are motile and tilted towards the caudal end (Thouvenin *et al*., 2020), we observed that motile cilia in 3 dpf larvae are located both on the ventral and dorsal wall of the central canal at the larval stage (Figure 4B). When we took a closer look at isolated motile cilia labeled in the *Tg(β-actin:Arll3b-GFP*) larvae at 3 dpf that were beating in the sagittal plane of imaging (see Methods), we observed that numerous motile cilia brushed against the RF in sweeping motions (Figures 4C1 and 4D1; see also star symbol in Movie 9), while others appear to almost stick to the RF with a glob of fluorescent material at their tip (Figures 4C2 and 4D2; see also Movie 9). We quantified the orientation of cilia in an angle relative to the horizontal (Figures 4E and 4F), and observed an overall tilt towards the caudal end both for dorsal inserted cilia (mean ± standard deviation provided hereafter: 64.8° ± 44.4°; N=49 cilia across 8 fish) and ventral inserted cilia (43.9° ± 35.8°; N=27 cilia across 9 fish; Figure 4G). We estimated the main ciliary beating frequency to be on average 11.2 Hz ± 2.4 Hz overall (Figure 4H; dorsal: 11.8 Hz ± 2.7 Hz; ventral: 11.6 Hz ± 2.8 Hz). However, we were limited in our acquisition frequency (40 Hz) in the setup that was suitable for ablation with a pulsed laser, thus the range of main beating frequencies up to only 20 Hz for both dorsal and ventral motile cilia may not reflect the actual beating frequency of these cilia (Figure 4H).

To investigate the impact of the RF on the beat frequency of motile cilia in the central canal, we performed acute focal ablations of the RF (see STAR Methods). We quantified the orientation and main beating frequency of the cilia before and after RF photoablation (Figure 4H; see also Supplementary Figure 1A and Movie 10). The tilt of cilia was not significantly different after RF photoablation (mean ± standard deviation provided hereafter: 55.1° ± 39.3°) from that before RF photoablation (57.4° ± 42.5°), illustrating that cilia orientation was, on average, not significantly affected by RF photoablation (N = 76 cilia from 9 fish; paired two-tailed t-test: p > 0.3) (Figure 4J). In contrast, the impact of acute RF photoablation on ciliary beating frequency showed different responses among individuals (Supplementary Figures 1B1-1B3). The fact that motile cilia densely surround the fiber along the walls of the central canal and brush along the fiber with a caudal tilt suggest that cilia-fiber interaction may generate friction promoting the oscillatory deflections of the RF observed *in vivo* (Figure 1) and of the deformation of the “flower” ends during RF relaxation after ablation (Figures 3B, 3B1, 3B2). Furthermore, perhaps these weak interactions observed between the fiber and beating cilia could explain heterogenous and graded tension we observed on the Reissner fiber, with larger tension on the rostral side.

**Figure 4.**
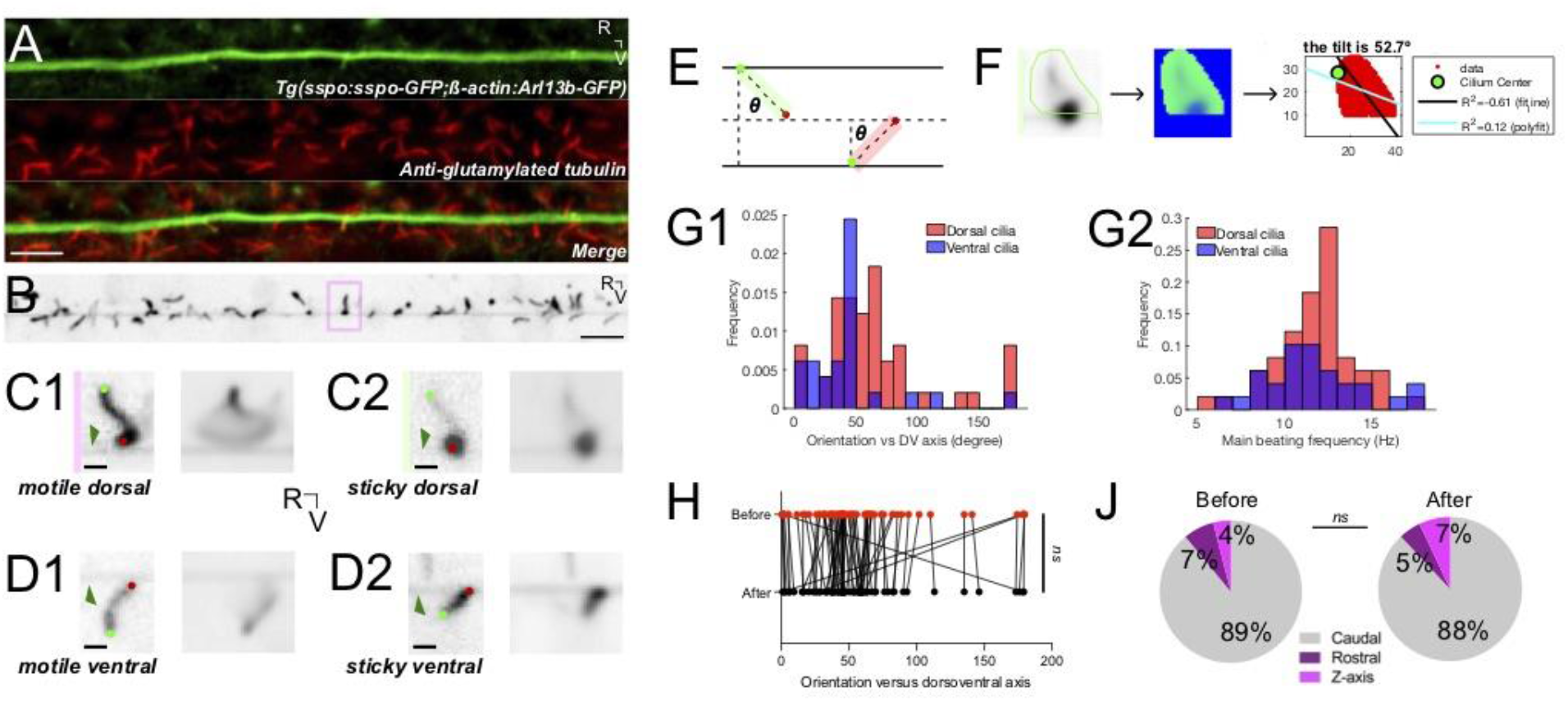
The Reissner fiber interacts with beating cilia protruding in the central canal. (A) Z-stack projection in the sagittal plane showing the immunostaining against GFP and glutamylated tubulin in a *Tg*(*sspo:sspo-GFP;β-actin:Arl13b-GFP*) larva fixed at 3 dpf revealing respectively the RF and cilia labeled by the *β-actin* promoter (both in green here) and motile cilia (red). Rostral to the left, ventral displayed on the bottom. (B) Single optical section in the sagittal plane showing the Reissner fiber surrounded by cilia protruding in the central canal in 3 dpf *Tg*(*sspo:sspo-GFP;β-actin:Arl13b-GFP*) paralyzed larva out of the streaming video acquired at 40Hz using spinning disk confocal microscopy setup equipped with a 40X objective. (C1) Right: example single optical section of a dorsal motile cilium from the highlighted box in (B). Dark green arrow indicates the RF. Light green and red dots indicate the base and tip of the cilium, respectively. Left: average projection over 25 s of the dorsal cilium brushing against the RF. (C2) Right: example single optical section of a dorsal motile cilium whose tip tends to stick to the RF. Left: average projection over 25 s of the dorsal cilium. (D1) Right: example single optical section of a ventral motile cilium whose tip tends to brush against the RF. Left: average projection over 25 s of the ventral cilium. (D2) Right: example single optical section of a ventral motile cilium whose tip tends to stick to the RF. Left: average projection over 25 s of the ventral cilium. (E) Schematic illustrating the geometric calculation of ciliary orientation versus the dorsoventral axis. The red rectangle represents a caudally-oriented ventral cilium, with 0° representing a completely caudally-oriented cilium and 90° representing the vertical with no tilt. The green rectangle represents a caudally-oriented dorsal cilium. For dorsal and ventral cilia, the caudal tilt corresponds to values between −90° and 90° with values greater than 90° corresponding to a rostral tilt. (F) Representative analysis process for dorsal cilium in (C2), with the dotted line representing the dorsal wall of the central canal and the green and red dots indicating the base and tip of the cilium, respectively. The green arrow indicates the RF in the middle of the central canal. A mask was first drawn over a single cilium to calculate the frequency and orientation in a specific region of interest (left). A temporal mean of that region (middle) was then used to calculate the orientation of the cilium in respect to the dorsoventral axis (right; see also Methods). (G) Distribution of the orientation in respect to the dorsoventral axis of dorsal (mean ± standard deviation provided hereafter: 64.8° ± 44.4°; 49 cilia across 8 fish) and ventral (43.9° ± 35.8°; 27 cilia across 9 fish) motile cilia. The dotted line indicates 90°, where values above 90° correspond to a rostral tilt. (H) Distribution of main ciliary beating frequency for dorsal (mean ± standard deviation provided hereafter: 11.8 Hz ± 2.7 Hz; 49 cilia) and ventral (11.6 Hz ± 2.8 Hz; 27 cilia) motile cilia. (I) The orientation of motile cilia in the central canal was not significantly different after RF photoablation (mean ± standard deviation provided hereafter: 55.1° ± 39.3°) from that before RF photoablation (57.4° ± 42.5°), illustrating that cilia orientation is, on average, not significantly affected by RF photoablation (76 cilia from 9 fish; paired two-tailed t-test: p < 0.3). The dotted line indicates 90°. (J) Overall rostrocaudal polarity of motile cilia remained similar before and after RF photoablation, with about 90% of motile cilia polarized toward the caudal end of the fish, and the remaining 10% either polarized toward the dorsal end of the fish or beating in the Z axis (76 cilia from 9 fish; paired two-tailed t-test: p < 0.3). ns = not significant Scale bar is 10 μm (A, B) and 2 μm (C1, C2, D1, D2).

## DISCUSSION

Our work reveals that the Reissner fiber is a dynamic structure under tension in the central canal *in vivo* with elastic properties and spontaneous oscillatory activity. Our mechanical model reveals that the Reissner fiber in larval zebrafish can be described as a soft elastic polymer that is maintained under tension in the CSF and exhibits a mechanical diffusion coefficient of 5 mm^2^/s, a characteristic time in the order of 2 ns and an elastic modulus of 10^6^ Pascal, which would fit more with the fiber acting as a low density gel rather than a rigid proteinaceous fiber. At baseline, in paralyzed and straight animals, we found evidence that the Reissner fiber interacts with the numerous long beating monocilia of ependymal radial glial cells (ERGs, which we estimated based on their density to at least hundreds of thousands at this stage), as well as with some ciliated sensory neurons (CSF-cNs, known to be only about a hundred at this stage, Prendergast *et al., in press*), whose activity decreases upon photoablation of the fiber.

### The Reissner fiber under tension oscillates along the dorsoventral axis in the central canal

By investigating the dynamical properties of the Reissner fiber in the central canal, we observed that the fiber is under tension *in vivo*, comforting previous observations of the RF being rectilinear (Troutwine *et al*., 2020) and in contrast with observations in the tissue after fixation, in which it curls and bends across the rostrocaudal axis (Orts-Del’Immagine *et al*., 2020). Our observations demonstrate that the RF can be modeled as a taut polymer under a heterogeneous tension *in vivo*. In addition to the spontaneous slow translation of the RF previously-observed along the rostrocaudal axis with material continually added and retracted from its surface (Troutwine *et al*., 2020), our observations further reveal spontaneous oscillatory activity of the fiber over the dorsoventral axis. We observe large spontaneous oscillatory activity of the RF in the middle portion of the fish, away from the attachment points in the rostral end (SCO) and caudal end (*ampulla caudalis*) (O. Meiniel & A. Meiniel, 2007; Gobron *et al*., 2000), akin to oscillatory behaviors of a plucked guitar string.

To capture enough photons from the GFP-tagged fiber in our recordings, we sampled displacements of the fiber along the dorsoventral axis at 40 Hz at most. However, we have indications that the dorsoventral oscillations of the fiber probably occur at higher frequency than the 20Hz we could observe in these conditions. Previously, the coordinated beating of motile cilia in the central canal has been reported up to 45 Hz, generating CSF flow estimated at a velocity of 10 *μm/s* in the embryo (Thouvenin *et al*., 2020). Such beating of cilia may influence locally how the Reissner fiber oscillates across the dorsoventral axis. Furthermore, interactions between motile cilia and the fiber may contribute to maintaining the fiber under tension with a graded tension on the rostral end. Further investigations of the interaction between the Reissner fiber and motile cilia in the CSF will be necessary to better understand the role of these interactions for the dynamic and physical properties of the RF.

### Elastic properties of the Reissner fiber subject to a heterogeneous tension decreasing along the rostrocaudal axis

Acute focal ablation of the RF allowed for the estimation of its elastic properties in the central canal. Upon ablation, the fiber retracts both further and faster when the ablation occurs on the rostral side of the fish compared to when the fiber is cut in the middle or caudal portions. A simple model of the RF represented as an elastic polymer under tension enabled us to estimate RF mechanical diffusion coefficient D ~10 mm^2^ /s and RF characteristic mechanical time **τ** ~10^-9^s. The full retraction time after the RF rupture is proportional to **τ** and to a huge factor (L / r_f_)^2^, with r_f_ of ~100 nm and L = 1mm. The value of the stress diffusion coefficient D being highly conserved in the rostral and middle/caudal portion of the fiber suggests that the diameter of the fiber is constant along the rostrocaudal axis. In contrast, the faster retraction speeds observed after ablation in the rostral side reveal that the pulling force on the fiber may not be uniform along the rostrocaudal axis. A higher tension in the rostral portion may be due to external factors increasing tension on the rostral side, such as the CSF flow going in the rostrocaudal direction (Tumani *et al*., 2017; Zhang *et al*., 2018)and/or the friction associated with the numerous interactions occurring between the fiber and the beating cilia along the central canal.

### Interactions between the Reissner fiber and ciliated cells in the central canal

We observed in streaming acquisitions of the RF and cilia labeled in GFP that some motile cilia brush and interact with the fiber. Accordingly, we measured that motile cilia from different fish have different beating frequencies in response to RF photoablation. Altogether, our observations indicate that RF interacts with motile cilia, a mechanism that can lead to reduction of ciliary beating as well as to friction between RF and the cilia. The impact of RF ablation on cilia beating frequency is still elusive, which is consistent with our previous observations that the absence of a fiber did not lead to a massive change in particle velocity profile in the embryo (Cantaut-Belarif *et al*., 2018). However, the fiber could nonetheless slightly alter the CSF flow profile in a transverse section by adding a point of null flow at the center of the central canal. Adding a point of null velocity can increase the velocity gradient in the CSF, and thereby increase the phenomenon of shearing at the level of the CSF-cN apical extension - in line with their functional coupling as reported was critical to mediate mechanoreception *in vivo* (Orts-Del’Immagine *et al*., 2020).

Since the Reissner fiber radius is ~ 100 nm (Orts-Del’Immagine *et al*., 2020), our resolution to identify the center of the fiber with classical fluorescent microscopy is certainly limited. Similarly, the precise mechanism of interaction between the RF and the ciliated neurons (CSF-cNs) cannot be finely resolved to decipher between direct contacts and a change of CSF flow at the level of the apical extension of the CSF-cNs. Nonetheless, we showed here that acute ablation of the Reissner fiber reduced the spontaneous calcium activity of CSF-cNs, indicating that a functional coupling may exist as well when the fiber spontaneously oscillates. Similarly, our analysis of ciliary beating was limited by the temporal resolution of our recordings on the confocal microscope equipped with a pulsed laser for ablation. A finer analysis of the spatiotemporal kinematics of the ciliary beating should resolve whether the Reissner fiber interacts preferentially with cilia beating at a given frequency, a given orientation or insertion in the central canal.

Here, we made a new step in understanding the axial sensory system composed of the Reissner fiber and the CSF-cNs. We demonstrated that the Reissner fiber at rest, when the body is straight, spontaneously oscillates and enhances spontaneous calcium activity in ventral CSF-cNs. The fiber’s vertical movement along the dorsoventral axis could lead to contact with CSF-cN apical extension to enhance the spontaneous calcium activity of ventrally-located sensory neurons we measured here. Interactions with dorsally-located sensory neurons may occur as well due to displacement in the horizontal plane, but were more difficult to image and report on in our conditions. The axial sensory system could form around the fiber showing oscillations and leading to the observed noise in the neuronal activity in resting baseline position (Sternberg *et al*., 2018; Prendergast *et al., in press*).

In zebrafish with genetic mutations, silencing or acute manipulations affecting CSF-cN activity or the maintenance of the Reissner fiber, sensorimotor deficits included slowing down of locomotion (a reduction of locomotor frequency and amplitude, Böhm *et al*., 2016; Wu *et al*., 2021),postural defects during fast locomotion (rolling, Hubbard *et al*., 2016; Wu *et al*., 2021) as well as morphological defects of the body axis in the embryo (Cantaut-Belarif *et al*., 2018) and the spine in juvenile/adult fish (Sternberg *et al*.,2018; Troutwine *et al*., 2020, Rose *et al*., 2020). Multiple evidence in mice indicate that CSF-cN-dependent functions for sensorimotor integration during locomotion observed in zebrafish are conserved in mammals (Gerstmann *et al*., 2022; Nakamura *et al*., 2022). The SCOspondin gene SSPOP is a pseudogene in humans (https://www-ncbi-nlm-nih-gov.proxy.insermbiblio.inist.fr/gene/23145), supporting the hypothesis that the Reissner fiber is absent in humans and great apes. However, evidence of urotensin signaling being relevant in human patients with adolescent idiopathic scoliosis (Dai *et al*., 2021; Xie *et al., in press*) raises the question of whether the urotensin signaling pathway involve CSF-cNs in order to contribute to morphogenesis and body axis straightening. Future studies will address whether acute ablation of the Reissner fiber can lead to similar deficits in sensorimotor integration for locomotion, posture and morphogenesis.

## Acknowledgements

We thank Sophie Nunes-Figueiredo, Monica Dicu and Antoine Arneau for fish care, the ICM Quant imaging facility and Xavier Baudin in the imaging facility of Institut Jacques-Monod for instrument use, scientific and technical assistance. We gratefully thank Prof. Brian Ciruna for the *Tg*(*β-actin:Arl13b-GFP*) transgenic line. This work benefited from equipment and services from the core facilities at the ICM (Institut du Cerveau, Hôpital Pitié-Salpêtrière, Paris, France). We thank all members of the Wyart lab for critical feedback (https://www.wyartlab.org). This work was supported by the HFSP Program Grants #RG0063 coordinated by Claire Wyart, in collaboration with Maria Lehtinen, Harvard University and François Gallaire, EPFL.

## Author contributions

CB performed all experiments and analyses in the manuscript. GZ & YCB made the first observations and estimations of the spontaneous translations of the Reissner fiber in live animals. YCB advised CB for imaging the Reissner fiber *in vivo* and performing IHC experiments. LM with discussions and tests from CB and CW generated the MATLAB scripts to locate the center position of the Reissner fiber along the rostrocaudal axis and estimate displacement in the dorsoventral axis as well as the PCA analysis script run on all pixels of the time series to estimate the major translation directions of the fiber. CW generated the MATLAB script to analyze the tilt and beating frequency of selected cilia beating in the sagittal plane of imaging. FBW and JFJ generated the model based on discussions with CB and CW, and provided valuable feedback to display the retraction kinematic after ablation. RSG provided the *Tg*(*sspo:sspo-GFP*) transgenic line that was instrumental for the observations on the dynamic of the fiber. CB & CW conceived the design of the ablation experiments using UV and infrared pulsed lasers, conceived all figures, and wrote the manuscript with inputs from all authors.

## SUPPLEMENTAL FIGURES

**Supplemental Figure 1.**
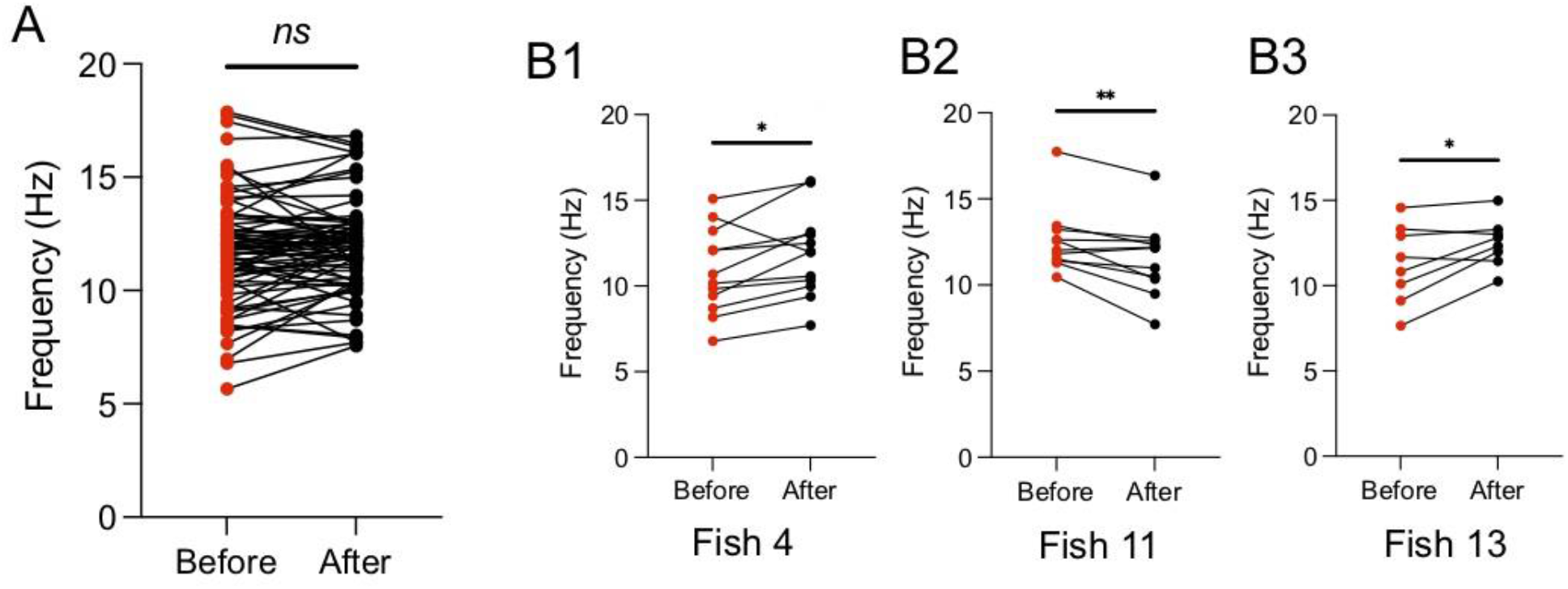
Changes in the main ciliary beating frequency varied from fish to fish in response to RF photoablation. (A) The main beating frequencies of dorsal and ventral motile cilia after RF photoablation (mean ± standard deviation provided hereafter: 11.9 Hz ± 2.0 Hz) compared to those before RF photoablation (11.7 Hz ± 2.4 Hz), illustrating that cilia main beating frequency was on average not significantly affected by RF photoablation (N=76 cilia from 9 fish; paired two-tailed t-test: p=0.3). (B1-B3) Select examples from (A), depicting fish whose response to RF photoablation significantly changed the main ciliary beating frequencies but in different ways. B1 and B3 illustrate an increase in main ciliary beating frequency after RF photoablation (B1: mean ± standard deviation provided hereafter: 11.9 Hz ± 2.5 Hz; B3: 12.5 Hz ± 1.4 Hz) from that before RF photoablation (B1: 10.9 Hz ± 2.5 Hz; N=12 cilia; paired two-tailed t-test: p=0.02; B3: 11.3 Hz ± 2.3 Hz; N=8 cilia; paired two-tailed t-test: p=0.03). B2 illustrates a decrease in main ciliary beating frequency after RF photoablation (11.6 Hz ± 2.2 Hz) from that before RF photoablation (12.6 Hz ± 1.9 Hz; N=11 cilia; paired two-tailed t-test: p=0.008).

**Supplemental Figure 2.**
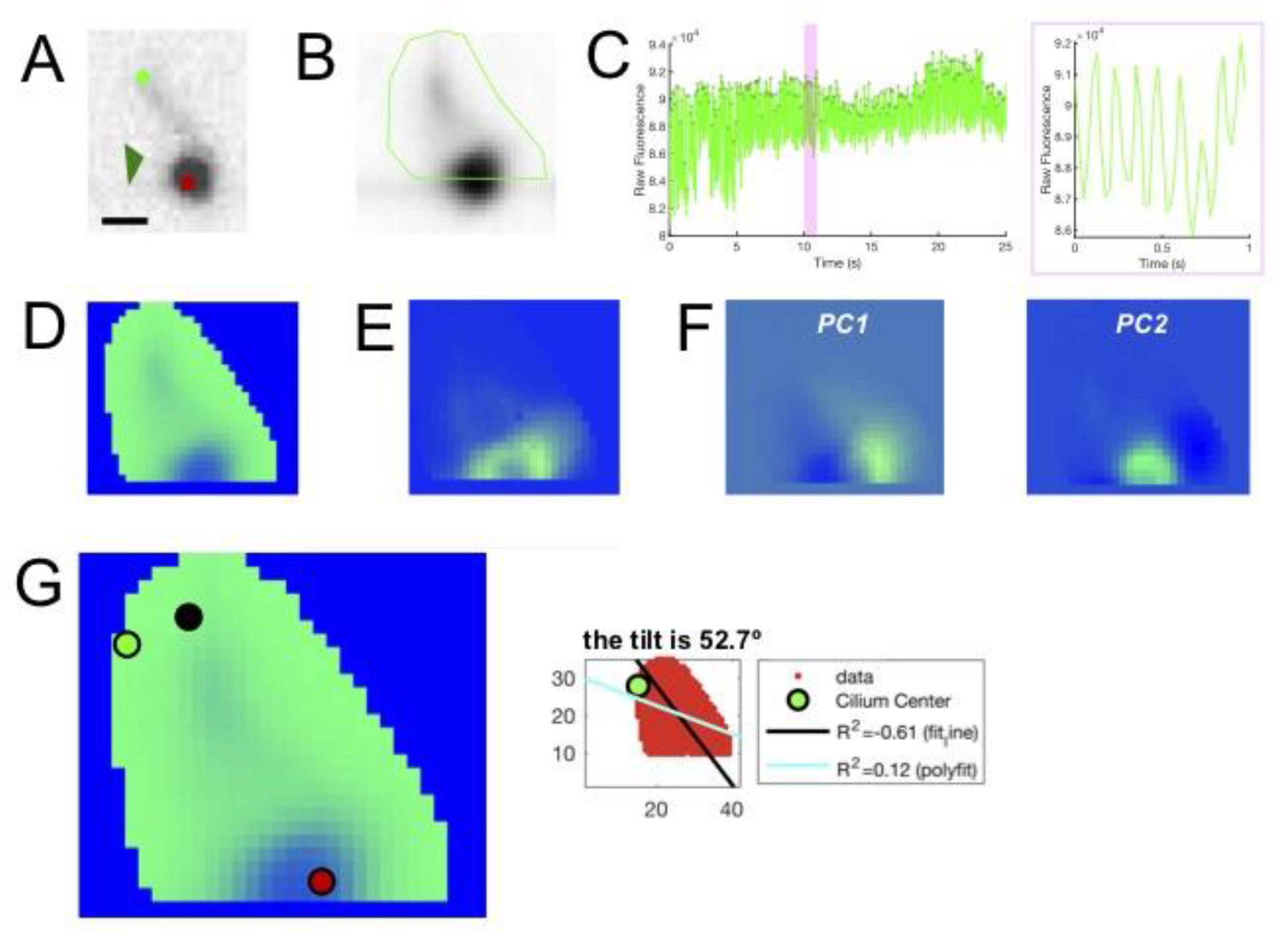
Example cilia analysis process. (A) Example of a single optical section showing a dorsal cilium from Figure 4C2. Dark green arrow indicates the RF. Light green and red dots indicate the base and tip of the cilium, respectively. (B) A mask drawn over the single cilium to calculate only the frequency and orientation in a specific region of interest. (C) The raw fluorescence trace of the inside of the mask over 25 s movie (left) and zoomed in over 1 s (right), corresponding to the highlighted area on the left panel, was used to detect peaks and estimate the ciliary beating frequency by counting the number of oscillations per second. (D) The temporal mean of the time series was calculated to see a temporally smoothed overview of the movie. (E) Standard deviation projection over 25 s before temporal smoothing. (F) A principal component analysis was performed to grasp which movements of the cilium could explain the most variability, which appeared in PC1 (left) and PC2 (right). (G) Ciliary orientation calculation. Left: window to calculate the orientation manually. Green dot represents the predetermined cilium center from the polyfit function in MATLAB. Black and red dots represent the manual inputs for the base and tip of the cilium, respectively. Right: The value for the manual calculation of orientation was compared to the orientation output of the two MATLAB functions and the one determined to best explain the data (in this case, a tilt of 52.7°) was kept for further analyses. Scale bar is 2 μm (A).

## STAR★METHODS

### Key Resources Table

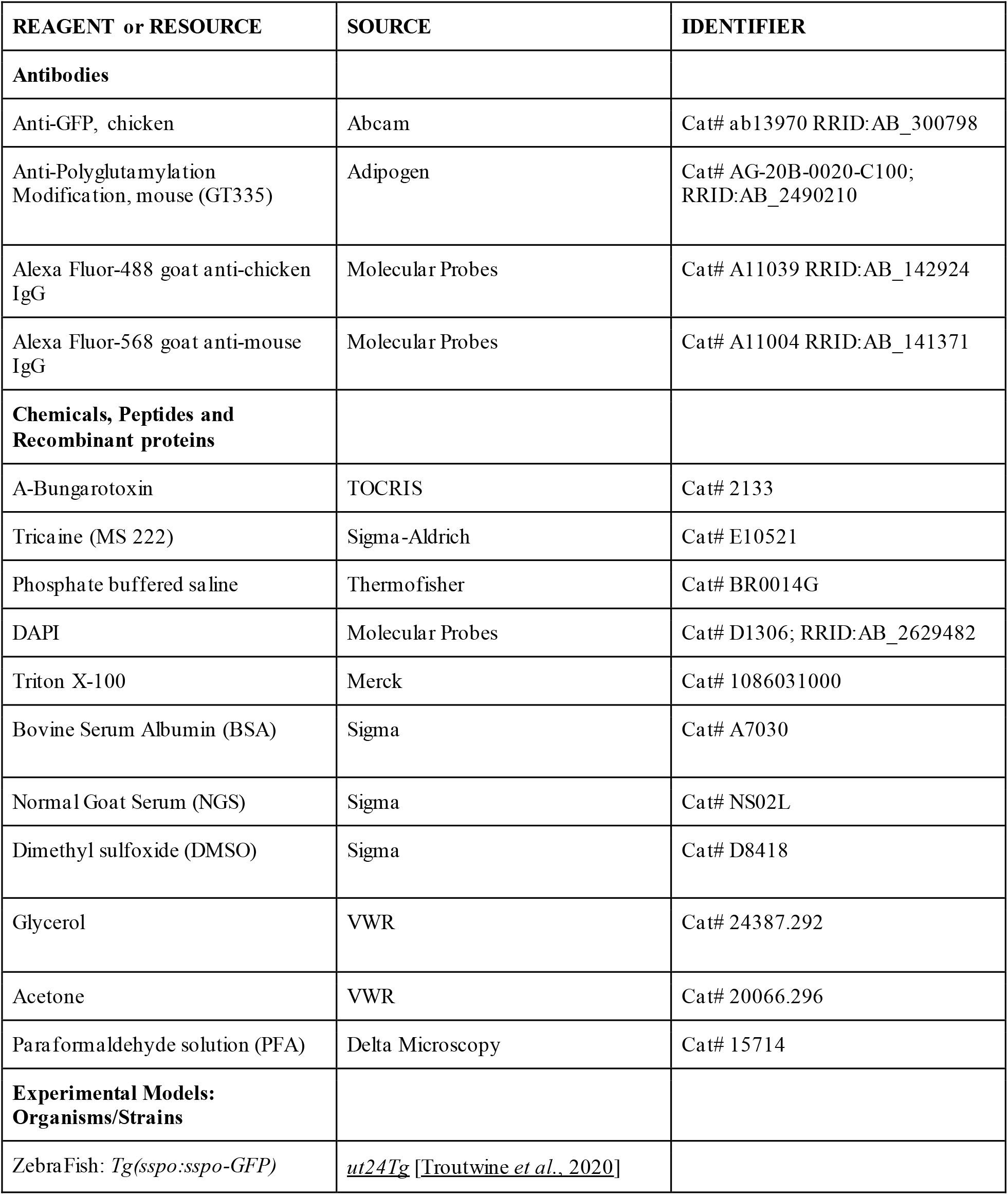
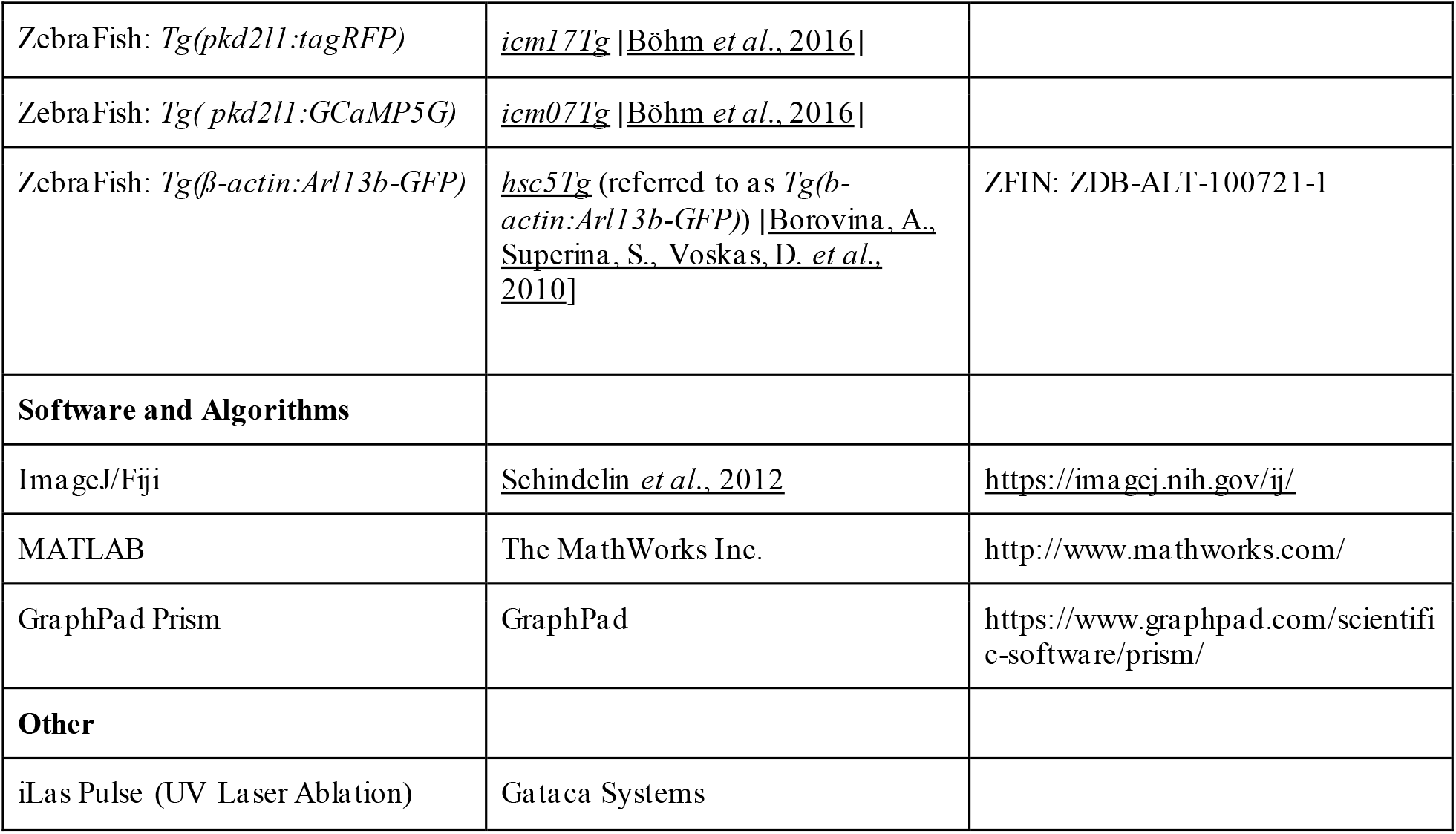

### Materials Availability Statement

Further information and requests for resources and reagents should be directed to and will be fulfilled by the corresponding author Claire Wyart. Note that this study did not generate new unique reagents. All data and codes generated and analyzed during this study can be found on Dryad (https://doi.org/10.5061/dryad.573n5tbc2).

### Experimental Model and Subject Details

All imaging procedures were performed on 3 days post fertilization (dpf) zebrafish larvae in accordance with the European Communities Council Directive (2010/63/EU) and French law (87/848) and approved by the Institut du Cerveau (ICM). All experiments were performed on *Danio rerio* embryos of AB, Tüpfel long fin (TL) and nacre background. Animals were raised at 28.5°C under a 14 / 10 light / dark cycle until the start of the experiment. All analyses were performed on animals that were in good health during the experiments.

### Method Details

All imaging experiments were done on 3 dpf *Tg*(*sspo:sspo-GFP*), *Tg*(*sspo:sspo-GFP;pkd2l1:tagRFP;pkd2l1:GCaMP5G*), and *Tg*(*sspo:sspo-GFP;β-actin:Arl13b-GFP*) zebrafish larvae. Their respective siblings were used as controls (i.e., wild-type and heterozygous mutants from the same clutch). For all experiments, larvae were laterally mounted in glass-bottom dishes (MatTek, Ashland, Massachusetts, USA), filled with 1.5% low-melting point agarose. Unless otherwise noted, larvae were paralyzed by injecting 1-2 nL of 500 mM alpha-bungarotoxin (TOCRIS) in the caudal muscles of the trunk via glass micropipette held by a micromanipulator (Märzhäuser Wetzlar MM-33), using a pneumatic Picopump (World Precision Instruments PV-820).

### Live imaging of the Reissner fiber

3 dpf larvae from *Tg*(*sspo:sspo-GFP*) incrosses were laterally mounted and paralyzed. An inverted spinning disk confocal microscope (Leica/Andor) equipped with a 40X water immersion objective (N. A. = 0.8) was used to acquire images at 40 Hz for 25 s. Fish were laterally sampled along the rostrocaudal axis from somites 1 to 30 to gain a holistic understanding of the Reissner fiber’s movements over the dorsoventral axis. The same procedure was performed with 3 dpf larvae from *Tg*(*sspo:sspo-GFP;β-actin:Arl13b-GFP*) double-transgenic larvae to image the Reissner fiber together with the motile cilia in the central to analyze how the Reissner fiber and motile cilia interacted in the central canal.

### Tracking of the Reissner fiber

To estimate the fiber curve *x↦y*(*x*) on each image, we implemented the following algorithm.

**Step 1:** First, we estimated the background level by analyzing the histogram of the whole sequence. Calling *I_0_* the intensity associated to the maximum value in the histogram (an approximation of the average background level), and *I_1_* the average of all intensities smaller than *I_0_* (so that *D*=*I_0_*-*I_1_* is an approximation of the average absolute deviation of the background), we subtracted the value *I_0_*+*D* to the whole sequence and set negative values to 0. This first step removes a lot of background noise from the sequence, without affecting much of the fiber signal. Then, we applied the following processing to each image of the sequence.
**Step 2:** We convoluted the image with a two-dimensional (non-isotropic) Gaussian filter to increase the signal-to-noise ratio. Since the fiber local orientation is always close to horizontal and has slow variations, we used a rather large horizontal width (*σ_x_* = 15/(3√2) ~ 3.5 pixels) and a moderate vertical width (*σ_y_* = 8/(3√2) ~ 1.9 pixel).
**Step 3:** For each column *x*, we computed the integer position *y_0_* of the pixel with maximal intensity *I(x,y_0_*), and used the intensities of the pixels above and underneath to get a sub-pixellic refinement *y_raw_(x*) of *y_0_* using a second order polynomial fit:

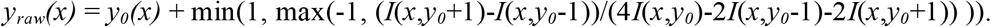
**Step 4:** At this stage, we also estimated the local fiber width as the diameter of the (one-dimensional) region obtained by thresholding the intensity at half the maximum value encountered on the current column. This value was used to classify columns as valid (domain *D*) when the width was less than 10 pixels, and invalid otherwise. This step is useful to remove uncertain position estimates due to firing neighboring cells: if both the fiber and a firing cell are visible (with similar intensity levels) on a given column *x*, the estimated width is too large and *x* will not belong to *D*.
**Step 5:** We then initialized *y(x*) to *y_raw_(x*) for all *x* and iterated until convergence a 2-step procedure:

1. *y*(*x*) ←y_raw_(*x*) for all *x* in *D* (enforce valid values for *y*)
2. *y*(*x*) ← (*y*\**G*)(*x*) /(1\**G*)(*x*) (smooth *y* using a convolution with a Gaussian kernel *G*). It is not difficult to prove that this iterative algorithm converges (independently of the initialization of *y*), and the convergence is quite fast. In practice, we use a Gaussian kernel *G* with σ~5.8, and 100 iterations were more than enough. Step 5 has two effects: it smoothes the function *y_raw_* and extrapolates it outside the domain *D*.
**Step 6:** In the (rare, but possible) case when there is a firing cell and an non-visible (or too faint) fiber on a given column *x*, Step 4 may wrongly classify *x* as a valid column. We can detect such a column *x* using the fact that the wrong estimate *y_raw_*(*x*) will generally be inconsistent with the expected fiber smoothness, and thus depart significantly from its corresponding smooth estimate *y*(*x*). In practice, we removed from set *D* all columns *x* for which |*y*(*x*)-*y*_raw_(*x*)| was larger than 1 pixel. With this restricted set *D*, we repeated the previous algorithm of Step 5 (iteration until convergence of the 2-steps procedure) to obtain the final estimate of the fiber curve *x↦y*(*x*).

The algorithm is made available as a MATLAB function. To prepare the acquired raw data to run the script, movies were cropped to the dimensions of roughly 15 μm x 100 μm, to have the Reissner fiber in the middle of the field of view with its surrounding background. Movies were discretized into 2 -μm bins along the rostrocaudal axis prior to running the script. When calculating dorsoventral displacement from the mean position of the fiber, movies were pieced apart and cropped to the dimensions of roughly 15 μm x 20 μm over 25 s five times, to encompass the whole recording cropped to 15 μm x 100 μm. The script was then run and dorsoventral displacement was saved to different variables and concatenated in the end to see the holistic dorsoventral displacement from the mean of the fiber for each fish.

### Two-photon ablation of the Reissner fiber and calcium imaging of ciliated sensory neurons

3 dpf larvae from *Tg*(*sspo:sspo-GFP;pkd2l1:tagRFP; pkd2l1:GCaMP5G*) incrosses were laterally mounted and paralyzed. Two-photon spiral scanning ablations at 800 nm were performed with a two-photon laser scanning microscope (2p-*vivo*, Intelligent Imaging Innovations, Inc., Denver, Colorado, USA) equipped with a 20X objective (N.A. = 1.0). A spiral scanning ablation at 800 nm over 0.5 μm was programmed within the two-photon photomanipulation settings before capturing the acquisition. Fish were laterally sampled using a 920 nm IR laser at an imaging frequency between 3 to 4 Hz over 75 s along the rostrocaudal axis from somites 1 to 30 to gain a holistic understanding of the impact of the RF ablation on CSF-cN activity before and after ablation along the central canal. Calcium transients of the CSF-cNs were analyzed by improving a previously-developed MATLAB function in order to reduce variability in the detection of the baseline in larvae compared to embryos (Sternberg *et al*., 2018). Movies were first renamed and renumbered in order to perform the analyses blindly. Movies were then registered to correct for any motion artifacts, and then run through the script to determine CSF-cN ROIs and their respective baselines for their calcium transient traces. CSF-cNs were only designated as active if there was a completed calcium transient greater than 3 standard deviations above the baseline over the 75 s recording.

### Ablation of the Reissner fiber with a UV pulsed laser for kinematic analysis of retraction

3 dpf larvae from *Tg*(*sspo:sspo-GFP;pkd2l1:tagRFP; pkd2l1:GCaMP5G*) incrosses were laterally mounted and paralyzed. An inverted CSU-X1 microscope (Yokogawa, Japan) equipped with a 40X oil immersion objective (N.A. = 1.3) was used to acquire images at 40 Hz for 25 s. A live FRAP photoablation (iLas Pulse, Gataca Systems, Massy, France) of the RF with an 8 nm pulse diameter and 20 ms duration was manually triggered in the middle of the acquisition. Prior to ablation, approximately 20 μm Z stacks (step size = 0.3 μm) were taken of the field of view at 10 Hz to gain a sense of the surroundings of the Reissner fiber at that position in the central canal. Fish were laterally sampled along the rostrocaudal axis from somites 1 to 30 to gain a holistic understanding of the Reissner fiber’s behavior upon ablation along the central canal. Subsequent analysis consisted of measuring the frame-by-frame relaxation of each end of the cut fiber, which was performed by a MATLAB function. Prior to running the script, movies were first rotated to have the Reissner fiber aligned horizontally at 0°, and then cropped to only include frames just before the UV ablation and until the fiber is out of the field of view.

### Ablation of the Reissner fiber with a UV pulsed laser with motile cilia imaging and analysis

A similar protocol as above was employed to ablate the Reissner fiber while observing the dynamics of the motile cilia. 3 dpf larvae from *Tg*(*sspo:sspo-GFP;β-actin:Arl13b-GFP*) double-transgenic larvae were laterally mounted and paralyzed. An inverted CSU-X1 microscope (Yokogawa, Japan) equipped with a 40X oil immersion objective (N.A. = 1.3) was used to acquire images for 25 s. Previous work at an imaging frequency of 100 Hz has shown main beating frequencies of motile cilia in the central canal ranging up to 45 Hz in 30 hours-post-fertilization zebrafish embryos (Thouvenin *et al*., 2020);however, in order to see the fluorescent signal of the RF at the larval stage, we performed these experiments at an imaging frequency of 40 Hz. A live FRAP photoablation (iLas Pulse, Gataca Systems, Massy, France) of the RF with an 8 nm pulse diameter and 20 ms duration was manually triggered in the middle of the acquisition. To analyze the resulting images, we implemented the following process:

**Step 1:** First, we selected individual dorsal and ventral motile cilia in sparse areas of each movie across 9 fish. Cilia that were chosen fulfilled the following criteria: 1) the individual cilium could be seen in focus (even if dimly fluorescent) during the duration of the movie, and 2) the cilium was in a sparse enough environment to analyze only its behavior. Cilia that were overlapping were therefore excluded from our analyses, since we could not distinguish distinct features from one cilium at a time if they were in this configuration. On average, approximately 9 individual cilia were selected in each recording and cropped in an approximately 10 μm x 10 μm square, to only get one specific cilium in the field of view.
**Step 2:** If after cropping another cilium or the Reissner fiber was still in the field of view, we would mask parts of the movie that had additional information via a MATLAB script. We drew a circle around the cilium of interest and masked the area outside the region of interest, to ultimately only run the analysis on the specific cilium of interest. The background of the movie was thresholded if the cilium was dimly fluorescent to increase signal and analysis accuracy.
**Step 3:** To calculate the ciliary beating frequency, the raw fluorescent trace of the signal inside a region of interest covering a motile section of the cilia was plotted. The main ciliary beating frequency in Hz was estimated by counting the number of oscillations in the fluorescent signal per second.
**Step 4:** The cilium’s orientation in respect to the dorsoventral axis was then calculated. Values from −90° to +90° relative to the horizontal with 0° toward the caudal end correspond to a caudal tilt of ventral cilia, while values from 0° to +90° degrees relative to the horizontal with 0° toward the caudal end correspond to a caudal tilt of dorsal cilia. Orientations of cilia were determined after comparing output values using two different MATLAB functions, in addition to a manual calculation, to conclude which value best fit the data.

Quantifications of cilia orientation in respect to the dorsoventral axis, cilia polarity direction and main ciliary beating frequency were then calculated for specific cilia before and after RF photoablation.

### Immunohistochemistry

Experiments were done on 3 dpf *Tg*(*sspo:sspo-GFP;β-actin:Arl13b-GFP*) double transgenic larvae. Larvae were first euthanized with 0.2% Tricaine and then fixed in a solution containing 4% paraformaldehyde solution (PFA), 1% DMSO and 0.3% Triton X-100 in PBS (0.3% PBSTx) at 4°C overnight. The samples were then washed once with 0.3% PBSTx to remove any traces of the fixation solution. For permeabilization, samples were incubated for 10 minutes at −20°C with acetone. Subsequently, samples were washed with 0.3% PBSTx (3×10 minutes) and blocked in a solution containing 0.1% BSA and 0.3% PBSTx for 2 h at room temperature. Samples were incubated overnight at 4°C with glutamylated tubulin (GT335, 1:400, Adipogen) for staining motile cilia, and GFP antibody (AB13970, 1:400, Abcam) to amplify the signals of the Reissner fiber and motile cilia in the primary antibody solution containing 0.1% BSA and 0.3% PBSTx. The next day, samples were washed with 0.3% PBSTx (3×1 h) and subsequently incubated overnight at 4°C with the secondary antibodies, Goat anti-Chicken Alexa Fluor Plus 488 (1:500) and Goat anti-Mouse Alexa Fluor 568 (1:500) (Thermofisher Scientific), with DAPI (1:1000) to stain for neuronal markers. After incubation with the secondary antibody, the larvae were washed (0.3% PBSTx, 3×1 h) and mounted on glass slides in Vectashield (Vector Laboratories, Burlingame, California, USA) to then be imaged using an inverted confocal SP8X White Light Laser Leica microscope with a 20X objective (N.A. = 1).

Immunohistochemistry experiments were also done on 3 dpf *Tg(sspo:sspo-GFP*) transgenic larvae to amplify the signal of the Reissner fiber. Larvae were first euthanized with 0.2% Tricaine and then fixed in a solution containing 4% paraformaldehyde solution (PFA) in PBS at 4°C overnight. The samples were then washed once with PBS to remove any traces of the fixation solution. Samples were blocked in a PBS solution containing 0.5% TritonX, 1% DMSO and 10% NGS at 4°C overnight. The next day, samples were incubated overnight at 4°C with GFP antibody (AB13970, 1:400, Abcam) in the primary antibody solution containing 0.5% TritonX, 1% DMSO and 1% NGS. On the third day, samples were washed with in a PBS solution containing 0.5% TritonX and 1 % DMSO (3×15 min), and subsequently incubated overnight at 4°C with the secondary antibody, Goat anti-Chicken Alexa Fluor Plus 488 (1:500) (Thermofisher Scientific), with DAPI (1: 1000) to stain for neuronal markers. After incubation with the secondary antibody, the larvae were washed in a PBS solution containing 0.5% TritonX and 1 % DMSO (3×15 min) and mounted on glass slides in Vectashield (Vector Laboratories, Burlingame, California, USA) to then be imaged using an inverted confocal SP8X White Light Laser Leica microscope with a 20X objective (N.A. = 1).

## SUPPLEMENTAL MOVIES

Movie 1a. **Time series showing the movement of the Reissner fiber in the sagittal plane of live *Tg(sspo:sspo-GFP*) transgenic larvae** (dorsal is up, rostral is left). Data was acquired at 40 Hz for 25 s.

Movie 1b. **Time series showing the lack of movement of the Reissner fiber in the sagittal plane in 3 dpf *Tg(sspo:sspo-GFP*) transgenic larvae <u>after fixation</u>** (dorsal is up, rostral is left). Data was acquired at 40 Hz for 25 s.

Movie 2. **Principal component analysis over all pixels of the time series reveal two main components of motion:** the first one (*top*) correspond to <u>translation</u> along the dorsoventral axis, and the second one (*bottom*) corresponds to <u>rotation</u>.

Movie 3a. **Spontaneous activity of cerebrospinal fluid-contacting neurons (CSF-cNs) in 3 dpf *Tg(sspo:sspo-GFP;pkd2l1:tagRFP; pkd2l1:GCaMP5G*) larvae before an acute ablation of the Reissner fiber.**

Movie 3b. **Spontaneous activity of cerebrospinal fluid-contacting neurons (CSF-cNs) in 3 dpf *Tg(sspo:sspo-GFP;pkd2l1:tagRFP; pkd2l1:GCaMP5G*) larvae after an acute ablation of the Reissner fiber.** Calcium imaging was recorded between 3 and 4 Hz over 75 s.

Movies 4-6. **Examples of retraction after laser ablation of the Reissner fiber used for figure 3 (A,B,C).** Position of the RF was tracked using a spinning disk operating at 40 Hz for 25 s.

Movie 7. **Motile cilia beating and interacting with the Reissner fiber in the *Tg(sspo:sspo-GFP;β-actin:Arl13b-GFP*) larvae.** Position of the cilia and RF were tracked using a spinning disk operating at 40 Hz for 25 s.

Movie 8. **Motile cilia beating after laser-mediated ablation of the Reissner fiber in the *Tg(sspo:sspo-GFP;β-actin:Arl13b-GFP*) larvae.** Position of the cilia were tracked using a spinning disk operating at 40 Hz for 25 s.

